# Chromosome Numbers and Reproductive Life Cycles in Green Plants: A phylo-transcriptomic perspective

**DOI:** 10.1101/2024.07.23.604804

**Authors:** Rijan R. Dhakal, Alex Harkess, Paul G. Wolf

## Abstract

The strong correlation between reproductive life cycle type and chromosome numbers in green plants has been a long-standing mystery in evolutionary biology. Within green plants, the derived condition of heterosporous reproduction has emerged from the ancestral condition of homospory in disparate locations on the phylogenetic tree at least 11 times, of which 3 lineages are extant. In all green plant lineages where heterospory has emerged, there has been a significant downsizing in chromosome numbers (refer to figure 1). This dynamic has been investigated without clear answers for many decades. In this study, we combine known ideas from existing literature with novel methods, tools, and data to generate fresh insights into an old question.

Using gene family evolution models and selection analyses, we identified gene families that have undergone significant expansion, contraction, or selection in heterosporous lineages. Our results reveal both shared and lineage-specific genomic changes associated with the evolution of heterospory. We found expansions in gene families related to developmental regulation, signaling pathways, and stress responses across heterosporous groups. Notably, the MATE efflux family showed consistent expansion and evidence of selection in heterosporous lineages, suggesting a potentially conserved role in heterospory evolution. These findings highlight novel insights that may underpin the association between heterospory and reduced chromosome numbers.

The general importance of chromosome numbers, structure, and sizes in cellular biology notwithstanding, the association between the emergence of heterosporous reproduction and chromosome number reduction/genome downsizing is not fully understood. It remains unclear why there exists an association between aspects of biology at such disparate levels as reproductive life-cycles and chromosome numbers/genome size. Exploring and answering this conundrum of evolutionary biology can add to our broader understanding of life sciences and of biology at different levels. Applying the novel tools and methods emerging from ongoing progress in biotechnology and computational sciences presents an opportunity to make new inroads into this long-standing question.

## Introduction

The evolution of land plants (Embryophyta) has been marked by several key innovations, including heterospory – the production of two distinct types of spores. This trait was a necessary precursor for seed-based reproduction, and has evolved independently multiple times across plant lineages. Notably, heterosporous plants, including all seed plants, generally have fewer chromosomes than their homosporous counterparts (Figure 1).

**Figure 1:-.**
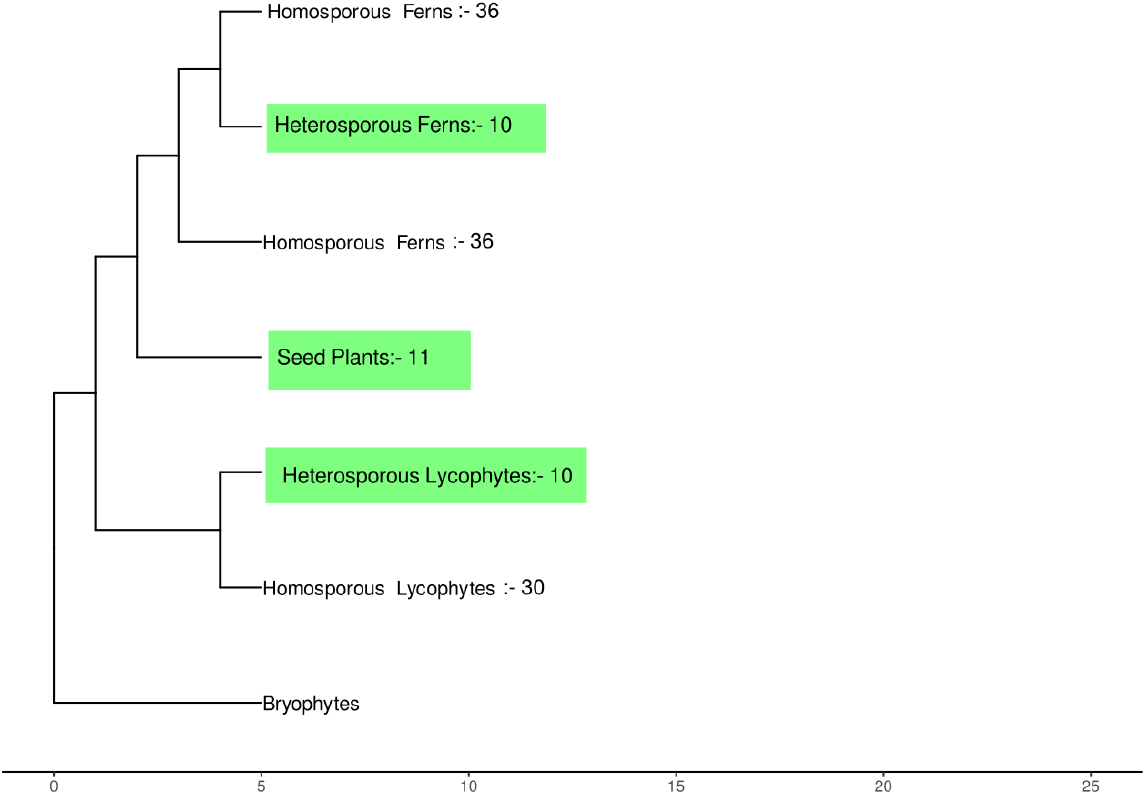
A simplified phylogeny of green plants/Embryophytes with the respective group and their average chromosome numbers in each group. Heterosporous lineages highlighted in green.

The pattern of reduced chromosome numbers in heterosporous lineages was first observed by (Klekowski and Baker 1966), who showed that homosporous plants have, on average, about four times as many chromosomes as heterosporous plants. However, the underlying mechanisms driving this association remain poorly understood. Previous hypotheses attempting to explain the higher chromosome numbers in homosporous lineages have been largely unsuccessful.

The gametophytic selfing hypothesis, proposed by (Hickok and Klekowski 1973), suggested that homosporous plants primarily reproduce by self-fertilization using gametes from a hermaphroditic gametophyte. This process would produce completely homozygous zygotes, leading to a complete loss of allelic diversity. The resulting loss of variation was hypothesized to create selective pressure favoring polyploidy-based redundancy to avoid genetic load, thereby leading to selection for larger genomes. However, this hypothesis was later rejected due to the tendency of polyploids to act as genetic diploids (C. H. Haufler and Soltis 1986). This diploidization process, where polyploid genomes undergo fractionation and gene loss to functionally resemble diploid genomes, would negate the proposed advantage of maintaining genetic diversity through polyploidy.

Recent genomic studies have provided new insights into this longstanding question. Contrary to earlier assumptions, homosporous plants appear to have lower rates of paleopolyploidy than heterosporous plants, despite their larger chromosome numbers. Instead, the high chromosome numbers in homosporous plants seem to result from higher retention of chromosomes from fewer rounds of polyploidy (Barker 2009; Christopher H. Haufler 2014; Marchant et al. 2021). This retention is likely due to lower rates of fractionation, i.e., the potentially permissive loss of duplicate genes and regulatory elements resulting from the relaxed effect of purifying selection on duplicated genomic elements.

These findings suggest a shift in perspective: rather than asking why homosporous lineages have larger chromosome numbers, we should investigate why heterosporous lineages have undergone greater genome downsizing (Kinosian, Rowe, and Wolf 2022). This reframing is particularly relevant given that heterospory is the derived condition within Embryophyta, having evolved independently at least 11 times across the plant evolutionary tree, including the three extant lineages.

The relationship between chromosome numbers and reproductive life cycles is not fully understood, despite its significance. Heterospory has a well-studied role as a necessary precursor to the emergence of seed bearing plants (Petersen and Burd 2017). Despite the crucial role of heterospory in the evolution of terrestrial plant life and its strong link to seed-based reproduction, the genomic mechanisms underlying transitions to heterospory and its association with reduced chromosome numbers remain unclear (Kinosian, Rowe, and Wolf 2022). Are there undiscovered patterns of neofunctionalization and/or subfunctionalization that affect reproduction and meiosis (and by extension chromosome numbers), shared among the extant heterosporous lineages?

Exploring the evolution of heterospory through a genomic lens can yield valuable new insights. However, within the phylogeny of embryophytes, a major challenge to advancing evolutionary biology is the scarcity of genomic data for some lineages. The phylogeny is replete with non-model plants lacking direct agronomic value or investment, resulting in limited data variety at the phylogenetic scale. In the absence of genomic data, transcriptomic data can serve as a valuable starting point. Recent initiatives, such as the One Thousand Plant Transcriptomes Project (One Thousand Plant Transcriptomes Initiative 2019), have significantly expanded the availability of transcriptomics data across diverse plant lineages. By combining these data with sophisticated phylogenetic and comparative genomic methods, including gene family evolution models and selection analyses, we now possess the tools to investigate patterns of genomic change associated with heterospory at unprecedented resolution and scale. This study leverages these modern data sources and methodologies to provide a fresh, data-driven exploration of the genomic underpinnings of heterospory and its relationship with chromosome number reduction.

In this study, we employ an exploratory approach to investigate potential genomic factors associated with the evolution of heterospory and its co-occurrence with reduced chromosome numbers. Our hypothesis is that similar genomic factors may be linked to both chromosome number reduction and the evolution of heterosporous lineages. We focus on two main questions:

- Have any gene families undergone significant expansion or contraction in copy number variations on lineages leading to heterosporous clades? If so, do their putative gene functions provide clues to the association between heterospory and chromosome number?
- Have any gene families undergone distinct and parallel patterns of selection on lineages leading to heterosporous clades? If so, do their putative gene functions provide insights into the association between heterospory and chromosome number?

By addressing these questions, we aim to provide new insights into the genomic basis of heterospory evolution and its potential link to chromosome number reduction. This investigation contributes to our understanding of plant evolution and has implications for agriculture and biotechnology, given the importance of seed plants in the human food supply chain.

To answer these questions, we employed a comprehensive genomic analysis approach, combining broad-scale examination with focused investigation of specific gene families. Our analysis pipeline involves sequence alignment, phylogenetic reconstruction, and statistical tests for gene family evolution and selection patterns. We applied this framework in two ways:

### Untargeted Approach

This novel approach aims to identify broad patterns of genomic change associated with heterospory across the plant phylogeny. Although this method offers the potential for unbiased discovery, we anticipate significant computational and algorithmic challenges due to the scale and depth of the data.

### Targeted Gene Family Analysis

To complement our genome-wide approach, we propose a targeted analysis focusing on specific gene families involved in cell cycling, spindle fiber formation, and chromatin dynamics – processes relevant to chromosomal profiles in a species. By narrowing our focus to these select gene families, we can mitigate several challenges:

- A reduced dataset allows for more intensive computational analysis, enabling the use of sophisticated algorithms impractical at a genome-wide scale.
- Focusing on specific gene families provides better phylogenetic resolution, especially for rapidly evolving genes problematic in genome-wide analyses.

Selecting genes with potential functional links to chromosome numbers increases the likelihood of detecting biologically meaningful patterns obscured in broader analyses. This targeted approach complements our untargeted analysis, allowing us to delve deeper into specific aspects of heterospory evolution while circumventing technical challenges inherent in large-scale genomic comparisons across diverse plant lineages.

Both approaches utilize a common set of bioinformatic tools and statistical methods, including sequence alignment, phylogenetic reconstruction, and tests for gene family evolution and selection patterns. By combining these complementary strategies, we aim to balance the potential for novel insights from the untargeted approach with the more directed examination of potential candidate genes.

## Materials and Methods

### Sample Selection

Our study aims to sample widely across land plants, encompassing all major clades of vascular plants and bryophyte outgroups. This comprehensive sampling is crucial for addressing our research questions about gene family evolution and selection patterns across heterosporous and homosporous lineages.

To achieve this goal, we balanced two competing factors: the need for a sufficiently large sample size to yield phylogenetic insights and the need for a manageable sample size to remain computationally feasible. Preliminary testing indicated that phylogenetic methodologies, including homology inference, gene family reconstruction, heuristics-based gene-tree inference, and selection analysis, remain tractable for up to approximately 100 species or transcriptomes using current software and hardware with realistic price tags.

To investigate specific alterations on heterosporous lineages, we included representatives from all three extant heterosporous lineages. We also sampled homosporous taxa evenly throughout lycophytes and ferns for evolutionary context. Additionally, we included outgroup samples from the three main bryophyte lineages. Our final dataset comprised 112 species, spanning at least 500 million years of evolutionary history(Wang et al. 2022).

### Data Sources

To investigate gene family evolution and selection patterns across plant lineages, we utilized genomic data from three different sources:

- (Pelosi et al. 2022): 4 species
- (Goodstein et al. 2012): 5 species
- One Thousand Plant Transcriptomes Project (One Thousand Plant Transcriptomes Initiative 2019): 103 species

We accessed data from Pelosi et al. (2022) from a public data repository (https://github.com/jessiepelosi/ferntxms). Phytozome data was accessed via their public website (https://phytozome-next.jgi.doe.gov/). We retrieved One Thousand Plant Transcriptomes Project data using an R package (https://github.com/ropensci/onekp), which enabled programmatic retrieval of OneKP data.

We used transcriptome assemblies as provided by the respective data sources, which had been quality-checked and filtered for low-quality reads prior to analysis.

This diverse dataset enables us to explore gene family expansions, contractions, and selection patterns across a wide range of plant lineages, directly addressing our research questions about the genomic basis of heterospory and its association with chromosome number reduction.

### Phylogenetic methodologies

#### Homology inference

To identify gene families shared among the studied species, a critical first step in addressing both of our research questions, we performed homology inference using OrthoFinder (v2.5.5) (David M. Emms and Kelly 2019) with default options. OrthoFinder was chosen for its ability to handle large datasets efficiently and its robust algorithm for identifying orthogroups. The bundled version of BLAST (McGinnis and Madden 2004) within the OrthoFinder installation was used as the sequence alignment tool. This generated a list of gene families (referred to as “orthogroups”) shared among the studied species. This homology inference step provides the foundation for our subsequent analyses of gene family evolution and selection patterns across the phylogeny.

#### De novo Species-tree generation

To provide a robust phylogenetic framework for our analyses of gene family evolution and selection patterns, we generated a species tree using STAG and STRIDE (D. M. Emms and Kelly 2018; David M. Emms and Kelly 2017). STAG constructs an unrooted species tree to account for multi-copy gene families, which STRIDE then roots. The final tree in Newick format is provided in the supplemental materials. This species tree is crucial for contextualizing the gene family expansions, contractions, and selection patterns we investigate in our study.

#### Gene family expansion and contraction inference

To address our first research question regarding gene family expansions and contractions in heterosporous lineages, we employed CAFE5 (Mendes et al. 2020), which employs a “birth-death” maximum likelihood model to reconstruct copy number variations at ancestral nodes for gene families. CAFE5 first estimates a global rate of change of evolution, or “lambda” and then employs that lambda in its implementation of maximum likelihood estimation.

Initially, we used CAFE5’s default settings to test the species tree generated by STAG and STRIDE, as well as all gene families identified by OrthoFinder. However, CAFE5’s inference model failed to initialize the exhaustive table of counts for all orthogroups. Through testing, we found that CAFE5’s statistical model fails when the difference between the largest and smallest copy number in a gene family exceeds 68. To overcome this limitation, we filtered out gene families exceeding this threshold, allowing us to proceed with the analysis while retaining the majority of gene families.

By reconstructing the history of gene family size changes across the phylogeny, this analysis allows us to identify gene families that have undergone significant expansion or contraction in heterosporous lineages, directly addressing our first research question.

#### Selection analysis

To address our second research question regarding distinct selection patterns in heterosporous lineages, we employed a multi-step approach:

1. Multiple sequence alignment: We performed multiple sequence alignment using both MUSCLE (Edgar 2004) and MAFFT(Katoh et al. 2002). These tools were chosen for their accuracy in handling large datasets and their ability to produce alignments suitable for downstream phylogenetic analyses. However, we encountered challenges with some of the largest gene families (in terms of sequence length and species diversity). For these families, the alignments produced had an excessive number of gaps. In fact, the gaps in these families were so prevalent that downstream codon alignment resulted in non-uniform length DNA sequences, which are incompatible with gene-tree inference methodologies. Consequently, the families that produced non-uniform sequence alignments were excluded from further analysis.
2. Translation to codons: We used PAL2NAL (Suyama, Torrents, and Bork 2006) to convert the aligned protein sequences to codon alignments. This step is crucial for maintaining the correct reading frame and for enabling subsequent analyses of selection at the codon level.
3. Gene tree generation: We generated gene trees for each gene family using IQ-tree2 (Minh et al. 2020) with the GTR+F+I model and 100 non-parametric bootstrap replicates. The GTR+F+I model was chosen as it allows for rate heterogeneity across sites and empirical base frequencies, which is appropriate for coding sequences and provides a robust framework for subsequent selection analyses.
4. Selection analysis: We employed the BUSTED-PH (BUSTED-PHenotype) method, an extension of the BUSTED (Branch-Site Unrestricted Statistical Test for Episodic Diversification) approach (Murrell et al. 2015). BUSTED-PH is specifically designed to test whether a particular feature, phenotype, or trait is associated with positive selection by comparing the selective pressures acting on gene families in lineages with and without the trait of interest.

The BUSTED-PH analysis requires a coding sequence alignment and a phylogenetic tree with the test and background branch sets clearly defined. We designated the branches leading to heterosporous lineages as the “test” set and the branches leading to homosporous lineages as the “background” set. Any remaining branches were designated as “nuisance” and excluded from the comparisons.

The analysis proceeds by fitting several random effects branch-site models to the data:

a. An “unrestricted” model that allows both the test and background branch sets to have their own independent ω (dN/dS) distributions.
b. A “constrained” model where ω ≤ 1 is enforced on the test branches. A likelihood ratio test between this model and the unrestricted model is used to determine if the test branches are subject to episodic diversifying selection.
c. A “constrained” model where ω ≤ 1 is enforced on the background branches. A likelihood ratio test between this model and the unrestricted model is used to determine if the background branches are subject to episodic diversifying selection.
d. A “shared” model where the ω distribution is constrained to be the same for test and background branches. A likelihood ratio test between this model and the unrestricted model is used to determine if the selective regimes differ between test and background branches.

The results of these likelihood ratio tests are then interpreted using a decision tree to provide guidance on whether episodic selection is associated with the presence of heterospory.By applying BUSTED-PH to our dataset, we aim to identify gene families that may have undergone distinct selection pressures in heterosporous lineages. This analysis directly addresses our second research question by revealing potential genomic signatures associated with the evolution of heterospory and its link to reduced chromosome numbers.

### Data processing and visualization

Data processing and reshaping were performed using custom R and Python scripts (see supplemental section for code repository). Visualization of phylogenetic trees and gene family evolution patterns was accomplished with the GGtree package in R (Yu 2020). Additional visualizations were created using ggplot2 in R and matplotlib in Python.

#### Targeted subset

Given the scale of the data and the depth of the phylogeny at hand, the untargeted approach was likely to encounter computational and analytical challenges. In anticipation of these challenges, we implemented a targeted analysis as a subset of our broader study. The targeted study is simply an implementation of the same methods and algorithms on the whole dataset, with the following distinctions:

1. Gene selection: We focused on a curated list of genes involved in cell-cycling, spindle fiber formation, and chromatin dynamics, derived from The Arabidopsis Information Resource (TAIR)(Rhee et al. 2003).
2. CAFE5 analysis: While most analytical steps remained identical for both approaches, we specifically re-ran CAFE5 for the targeted gene set to allow for a more focused examination of gene family evolution in these genes of interest.
3. Computational resources: The reduced gene set in the targeted approach allowed for more intensive analysis and manual curation where necessary, without the computational constraints of the full dataset.

This targeted approach complements our untargeted analysis, providing a more detailed examination of specific gene families hypothesized to play key roles in heterospory evolution and chromosome number reduction.

## Results

### Species Tree Topology

The species tree generated using STAG and STRIDE is presented in Figure 2. This de novo generated tree provides the phylogenetic context for all subsequent analyses of gene family evolution and selection patterns. Importantly, the topology of our tree is largely consistent with current understanding of plant phylogenetic relationships as established in recent literature (Wickett et al. 2014; Ruhfel et al. 2014; One Thousand Plant Transcriptomes Initiative 2019).

### Untargeted analysis

#### Gene Family Expansions and Contractions

CAFE5 revealed significant changes in gene family sizes across the phylogeny, particularly in lineages leading to heterosporous plants. We detected 54 gene families with significant expansion or contraction leading to the three heterosporous nodes in our phylogeny. The distribution of these changes varied across the heterosporous lineages.

**Table 1:-.** Counts of families that had experienced significant changes at the ancestral nodes that lead to heterospory as observed from an untargeted approach.

| Nodal Common ancestor | Families with significant quantitative change |
| --- | --- |
| Seed Plants | 12 |
| Heterosporous Ferns | 11 |
| Heterosporous Lycophytes | 4 |

To understand the functional implications of these changes, we examined the *Arabidopsis thaliana* (Rhee et al. 2003) orthologs within these gene families and their associated functions. Supplemental Tables 1-3 present the full list of gene families with significant changes for each heterosporous lineage. While all identified changes are potentially significant, several gene families stand out due to their known roles in plant development, reproduction, and stress responses:

#### Seed Plant common ancestor node

The ancestral node of seed plants exhibited expansions in gene families associated with reproductive adaptations. The MATE efflux family (OG0000126) showed significant growth, potentially enhancing control over flavonoid metabolism crucial for seed and pollen development. Concurrent expansions in LEA proteins (OG0000163) and seed imbibition-related genes (OG0000308) suggest a coordinated evolution of mechanisms supporting seed viability and germination efficiency.

#### Heterosporous Ferns common ancestor node

For the parent of heterosporous ferns, we observed a parallel expansion of the MATE efflux family (OG0000126), mirroring the pattern seen in seed plants. This convergence hints at a possible shared mechanism in heterospory evolution. The expansion of universal stress proteins (OG0000287) indicates enhanced stress resilience, while growth in the homeobox gene family (OG0000717) suggests changes in the developmental regulation pathways.

#### Heterosporous Lycophyte parent node

For the parent of heterosporous Lycophyte, the most prominent change was the expansion of homeobox genes (OG0000717). This growth implies an increased sophistication in gene regulation networks, potentially facilitating the complex life cycle transitions characteristic of heterosporous plants.

The dynamics of quantitative expansions/contractions across heterosporous lineages reveal a complex interplay of shared and lineage-specific genomic changes. We observed a consistent trend of expansions in gene families related to developmental regulation, signaling pathways, and stress responses across all three heterosporous groups. The recurrent expansion of certain families, such as MATE efflux and homeobox genes, across multiple lineages suggests potential common adaptive strategies in the evolution of heterospory. This pattern indicates that the transition to heterospory may be associated with increased genomic complexity in these functional categories, likely reflecting the need for more sophisticated regulation of reproductive processes and adaptations to new ecological niches. However, the unique expansion patterns in each group also highlight the diverse evolutionary paths to heterospory, underscoring the flexibility of genomic solutions to this reproductive innovation.

It is important to note that our analysis was limited to gene families with moderate levels of copy number variation due to computational constraints. Rapidly evolving gene families (CNV > 68) were excluded from this analysis, which may have led to the omission of some relevant genomic changes.

### Selection Analysis Results

Of the 54 gene families identified in our CAFE analysis, 51 were successfully analyzed using BUSTED-PH. Three families were excluded from this analysis due to challenges in producing uniform length sequence alignments, which are necessary for the BUSTED-PH method. This limitation highlights the computational and methodological challenges in analyzing rapidly evolving or highly divergent gene families.The detailed outcomes of BUSTED-PH are in supplemental tables 4 through 6.

### Selection Analysis Across Heterosporous Lineages

Our BUSTED-PH analysis revealed diverse patterns of selection across the three heterosporous lineages: angiosperms (representing seed plants), heterosporous ferns, and heterosporous lycophytes. In angiosperms, all 11 gene families with proper codon alignment showed some form of selection related to the heterospory phenotype. Specifically, three families (OG0000126, OG0000108, OG0000104) exhibited strong evidence of selection in the heterosporous lineage. Six families displayed different selection regimes between heterosporous and homosporous lineages, while two families (OG0000145, OG0000451) showed similar selection regimes across both lineage types. Notably, the MATE efflux family protein (OG0000126) demonstrated significant expansion and strong heterospory-specific selection, underscoring its potential crucial role in seed plant evolution.

In heterosporous ferns, analysis of eight gene families revealed similarly diverse selection patterns. Five families (OG0000126, OG0000717, OG0001114, OG0002601, OG0000860) showed strong evidence of heterospory-specific selection. Two families exhibited distinct selection regimes between heterosporous and homosporous lineages, while one family (OG0000857) showed selection in the heterosporous lineage without significant difference from homospory. Remarkably, the MATE efflux family protein (OG0000126) again exhibited both expansion and strong heterospory-specific selection, mirroring its pattern in angiosperms and suggesting a conserved role in heterospory evolution across plant lineages.

For heterosporous lycophytes, analysis of four gene families revealed a distinct selection pattern. Three families (OG0000022, OG0000171, OG0000598) exhibited selection in both heterosporous and homosporous lineages, but with different regimes. This suggests that while these gene families are under selection pressure in both lineage types, the nature of the selection differs between heterosporous and homosporous lycophytes. One family (OG0000634) could not be analyzed due to misaligned codon translation, an issue arising from significant evolutionary divergence. Interestingly, the gene family OG0000171, encoding transcription factors with JmjC domains (likely involved in histone demethylation), showed significant expansion in our CAFE analysis and evidence of differential selection between heterosporous and homosporous lineages.

Comparing selection patterns across the three heterosporous lineages reveals several intriguing trends. The MATE efflux family protein (OG0000126) consistently showed strong evidence of heterospory-specific selection in both angiosperms and ferns, indicating a potentially conserved role in heterospory evolution. However, many gene families exhibited different selection patterns across lineages, suggesting that the genomic basis of heterospory may have evolved independently in each group. While most gene families in all lineages showed evidence of selection, the regimes often differed between heterosporous and homosporous lineages. This suggests that the nature of selection pressure may have shifted with the evolution of heterospory.

These findings provide evidence for both shared and lineage-specific genomic changes associated with the evolution of heterospory. The combination of gene family expansion and distinct selection pressures in heterosporous lineages suggests that the transition to heterospory may have involved significant genomic restructuring and adaptation, although the specific mechanisms vary across plant groups.

### Targeted analysis results

#### Gene family expansion contractions

**Table 2:-.** Counts of families that had experienced significant changes at the ancestral nodes that lead to heterospory as observed from a targeted approach.

| Nodal Common ancestor | Families with significant quantitative change |
| --- | --- |
| Seed Plants | 0 |
| Heterosporous Ferns | 1 |
| Heterosporous Lycophytes | 1 |

When CAFE5 was applied to our targeted subset of genes, only one gene family showed significant expansion on ancestral nodes shared by certain heterosporous lineages. Specifically, the gene family OG0000171 exhibited expansion on all ancestral heterosporous nodes, except the one common to the angiosperms. In other words, it saw significant expansion on nodes ancestral to both heterosporous ferns and lycophytes.

Functionally, this gene family encodes transcription factors with JmjC domains. These proteins are known to function as histone demethylases, playing crucial roles in epigenetic regulation. Notably, OG0000171 was also identified in our untargeted CAFE analysis as showing significant expansion in the heterosporous lycophyte lineage. Furthermore, our untargeted selection analysis revealed evidence of differential selection between heterosporous and homosporous lineages for this gene family.

### Selection analysis

The results for the selection analysis of the list of targeted gene families in supplemental table 7 through 9.

### Selection Analysis Results for Targeted Gene Families

Our targeted analysis focused on gene families involved in cell cycling, chromatin dynamics, and spindle fiber formation - processes we suspect are potentially crucial to chromosome behavior and reproductive mechanisms. The BUSTED-PH analysis revealed a complex landscape of selection pressures acting on these genes in heterosporous lineages, providing insights into the genomic underpinnings of this major evolutionary transition.

Given the complex nature of heterospory as a reproductive strategy, we hypothesized that its evolution might involve myriad changes across multiple cellular processes. Our results indeed reveal a diverse array of selection patterns across the analyzed gene families, consistent with this hypothesis.

Key findings from our analysis include:

- Heterospory-Specific Selection: A substantial number of gene families across all three categories showed strong evidence of selection specifically in heterosporous lineages. This pattern was particularly pronounced in chromatin-related genes, with nearly 30% of families in this category exhibiting heterospory-specific selection.
- Differential Selection Regimes: Many gene families showed evidence of selection in both heterosporous and homosporous lineages, but with different regimes. This pattern was most common in chromatin-related genes.
- Conservation: A notable proportion of gene families across all categories showed no evidence of selection associated with heterospory.
- Subtle Shifts: Some gene families showed evidence of selection in heterosporous lineages, but without significant differences from homosporous lineages.

Among the three categories, chromatin-related genes showed the highest proportion of families under heterospory-specific selection, followed by spindle fiber genes. Cell cycling genes exhibited the most diverse range of selection patterns.The observed patterns suggest a complex interplay of genomic changes associated with this reproductive transition, involving both conservation of core functions and adaptive changes in others.

## Discussion

We identified significant expansions, contractions, and selection pressures in gene families on branches leading to angiosperms, heterosporous ferns, and heterosporous lycophytes. Our comprehensive analysis reveals a landscape of quantitative and qualitative evolutionary signals, showing both shared and unique changes in gene families of heterosporous lineages.

Our untargeted quantitative analysis revealed expansions in gene families related to developmental regulation, signaling pathways, and stress responses across all three heterosporous lineages. However, the specific families affected and the nature of selection pressures varied among lineages. Notably, many of these expanded families, particularly those involved in reproductive processes and hormone signaling, also showed evidence of differential selection.

Our targeted approach identified a gene family (OG0000171) encoding JmjC domain-containing proteins involved in histone demethylation, which showed significant expansion in heterosporous ferns and heterosporous lycophytes. This family was also identified in our untargeted analysis, highlighting the potential importance of epigenetic regulation in heterospory evolution. The expansion and strong selection pressure on the MATE efflux family protein (OG0000126) in both seed plants and heterosporous ferns suggest a potentially conserved role in heterospory evolution. Furthermore, the expansion of stress-response-related gene families, particularly in heterosporous lycophytes, may indicate adaptations to new ecological niches, facilitating the colonization of diverse terrestrial environments.

The MATE efflux family, showing both expansion and strong selection pressure in multiple heterosporous lineages, emerges as a particularly interesting candidate for further study. This finding aligns with established theories, such as the Haig and Westoby hypothesis (Haig and Westoby 1988), which posits that heterospory evolved due to increasing vegetative complexity and competitive pressures in early terrestrial environments. As conditions for the establishment of young sporophytes became more challenging, selection favored larger spores with greater nutrient reserves, leading to the evolution of a size threshold at which the production of small, male-only spores (microspores) alongside larger female spores (megaspores) became advantageous.

Our genomic findings offer new molecular insights into the Haig and Westoby model. The observed expansions in gene families associated with hormone signaling and stress responses across heterosporous lineages align with the ecological shifts proposed by Haig and Westoby. These genetic changes may have facilitated the adaptive responses necessary for survival in more complex and competitive environments. For example, the expansion of the MATE efflux family, involved in hormone transport and stress tolerance, may have played a crucial role in regulating the growth and development of both gametophytes and young sporophytes under challenging conditions.

The untargeted but context-aware selection analysis (BUSCO-PH) highlighted a high proportion of chromatin and spindle fiber genes under heterospory-specific selection. This suggests significant changes in gene regulation and cell division mechanisms during heterospory evolution, which could be critical for controlling gene expression during the development of distinct spore types, altering meiotic processes for megaspore and microspore production, and modifying chromosome behavior in gametophyte differentiation. The diverse selection patterns in cell cycling genes suggest a balance between innovation in reproductive strategies and the conservation of fundamental cellular processes.

The expansion and selection of chromatin-related genes in relation to the phenotype of heterospory underscore the potential role of epigenetic regulation in plant evolution and adaptation. As immobile organisms, plants require a high degree of plasticity to cope with their surrounding environmental factors. This finding is consistent with the increasing recognition of epigenetic mechanisms as crucial players in significant evolutionary and adaptive transitions, particularly the non-random nature of the inheritance of epigenetic adaptations (Miryeganeh and Saze 2020).

## Conclusions

We observed a complex interplay of quantitative and selective changes across various lineages, generating novel insights into the genomic mechanisms underlying the evolution of heterospory. The consistent expansion and selection of specific gene families, particularly those involved in developmental regulation, signaling pathways, and stress responses, suggest common adaptive strategies in heterospory evolution.

Although we did not detect direct genomic signals for chromosome number regulation, changes in developmental and epigenetic genes may indirectly influence meiotic processes or genome stability. This finding provides a new perspective on the long-standing question of reduced chromosome numbers in heterosporous plants. The differences in genomic changes among angiosperms, ferns, and lycophytes, despite their shared emergence of heterospory, reflect the flexibility of genomic pathways leading to this reproductive innovation.

The complex pattern of selection across gene families suggests that the transition to heterospory was a multifaceted evolutionary process involving changes in various cellular systems. This aligns with the multiple independent origins of heterospory in plant evolution, indicating both common genomic changes and flexibility in achieving heterospory.

Our study highlights the power and limitations of large-scale comparative genomic approaches. While we identified broad patterns across diverse plant lineages, computational constraints and reliance on model organism annotations posed significant challenges. These limitations underscore the need for more powerful computational tools and comprehensive functional annotations across non-model organisms.

Despite methodological constraints, our study has yielded insights into the genomic underpinnings of heterospory evolution. It demonstrates the potential of large-scale comparative genomics to address fundamental questions in plant evolution, while also revealing areas where methodological improvements are needed. Future studies will benefit from an interplay between targeted questions and more powerful algorithms, as well as increased computational power and expanded genomic data.

Advances in next-generation sequencing (NGS) and computational technologies will likely enhance the results of our experimental design in future iterations. Improvements in NGS technology will increase the availability of genomic data for non-model plants, while advancements in computational tools will enable the implementation of more sophisticated phylogenetic algorithms. This synergy between biological insight and methodological advancement will be crucial for deepening our understanding of major evolutionary transitions in plants and their implications for plant biology.

### Limitations of the Study

Our study has several limitations. One significant limitation is the nature of the data itself, as illustrated in supplemental Figures 1 and 2 with BUSCO (Benchmarking Universal Single-Copy Orthologs) scores (Simão et al. 2015). BUSCO is a metric used to assess the completeness of genome assemblies by evaluating the presence of a set of highly conserved genes expected to be found in nearly all eukaryotes.

These illustrations show that our data has a wide range of completeness and areas needing improvement. Notably, the absence of evidence does not necessarily imply evidence of absence. As seen in supplemental Figures 3 and 4, while the availability of non-model plant genomic data has improved significantly in recent years, further enhancement is still needed. The distribution of these scores highlights the progress made but also the challenges that remain in obtaining complete genomic data for diverse plant lineages. Additionally, relying on *Arabidopsis thaliana* for functional annotation may have introduced a bias towards angiosperm-centric functions. Our targeted approach, while focused, may have missed important gene families not included in our initial selection criteria.

We observed a discrepancy in the quantitative outcome for OG0000171 between our targeted and untargeted data processing approaches. In the untargeted CAFE5 analysis, OG0000171 was expanding only in heterosporous lycophytes, whereas in the targeted analysis, it significantly expanded in both heterosporous lycophytes and heterosporous ferns. This discrepancy highlights the differences between our targeted and untargeted approaches. The targeted approach allows for focused analysis of specific gene families, potentially increasing sensitivity to subtle changes. However, it depends heavily on existing functional annotations and prior hypotheses about relevant genes, which can be a limitation, especially when studying non-model organisms or evolutionary innovations with poorly understood genetics. In contrast, the untargeted approach offers a broader view of genomic changes, capable of identifying patterns across the genome, but its comprehensive scope can sometimes overwhelm the algorithm in CAFE5. Although both approaches found similar overall patterns, the untargeted approach showed attenuated results for certain gene families. This difference underscores the importance of cautious interpretation.

### Future Directions

This study leveraged contemporary data and methodologies to explore the longstanding mystery of heterospory in plant biology. We identified promising candidate genes for future experimental validation, particularly those exhibiting expansion and strong selection pressure. Further investigation into the functional characterization of these gene families and their roles in heterospory development and maintenance is warranted.

Ecological studies investigating the relationship between habitat characteristics—such as vegetative complexity and environmental stressors—and the degree of heterospory in extant species could provide empirical support for the Haig and Westoby model. These studies can be complemented by functional genomic experiments targeting MATE gene families, focusing on the impact of mutations in these families on *in-vivo* biology. Such experimentation would also enable testing for possible connections between MATE gene families and meiosis or chromosome numbers.

Future research should aim to bridge the gap between genomic insights and the ecological and developmental aspects of heterospory evolution. Investigating the expression patterns and functions of expanded gene families during spore and gametophyte development, particularly in relation to nutrient allocation and stress responses, would be beneficial. Comparative studies of spore development and early sporophyte establishment across species with varying degrees of heterospory could help elucidate how genetic changes translate into adaptive advantages in different environments. By integrating genomic findings with ecological and developmental approaches, we can advance our understanding of heterospory evolution. This comprehensive approach has the potential to clarify how genetic innovations facilitated major life cycle transitions proposed by Haig and Westoby, addressing longstanding questions about this pivotal innovation in plant evolutionary history.

Future research would benefit from the development of more sophisticated algorithms capable of handling rapidly evolving gene families without computational bottlenecks. This could include improvements to existing tools like CAFE5 or the creation of new tools better suited to handle extreme variations in gene family size across evolutionary time. Advancements in sequence alignment techniques, especially those sensitive to gene families evolving over longer time scales, could significantly enhance our ability to detect and analyze genomic changes associated with major evolutionary transitions like heterospory.

While integrating other types of data, such as metabolomics or detailed phenotypic information, could provide a more comprehensive picture of heterospory evolution, practical challenges in obtaining such data across diverse plant lineages are considerable. In the interim, comparative studies focusing on genomic regions surrounding the gene families discovered in this study could help investigate potential regulatory changes driving observed patterns of gene family evolution and selection.

## Supporting information

The de-novo species tree used through out the manuscript.

## Acknowledgements

The author would like to acknowledge the following individuals/organizations for their lending their time, resources and expertise:-

This project was funded by the U.S. National Science Foundation, NSF Grant #1911459 to PGW.

Alabama Supercomputing Authority:-This project had generous access to compute power thanks to the Alabama Supercomputing Authority.

Or helpful comments on earlier drafts of the manuscript:

Norm Wickett, PhD - https://orcid.org/0000-0003-0944-1956

Sylvia Kinosian, PhD - https://orcid.org/0000-0002-0918-7196

## Author Contributions

Rijan Dhakal :- Conducted the research and wrote the paper.

Alex Harkess :- Significant consultation, advisory and guidance on the methods and manuscript.

Paul Wolf:- Secured funding and critically supervised the research and drafting of the manuscript.

## Supplemental Data:-

1. The code that was used to conduct this research is available here:- https://github.com/VectorFrankenstein/spory-phylogenetics-project-code
2. The upstream data, intermediates and final figures and tables are hosted with Zenodo here:- https://zenodo.org/records/13895891

